# Genomic signatures of sympatric speciation with historical and contemporary gene flow in a tropical anthozoan

**DOI:** 10.1101/399360

**Authors:** Benjamin M. Titus, Paul D. Blischak, Marymegan Daly

## Abstract

Sympatric diversification is increasingly thought to have played an important role in the evolution of biodiversity around the globe. However, an *in situ* sympatric origin for co-distributed taxa is difficult to demonstrate empirically because different evolutionary processes can lead to similar biogeographic outcomes-especially in ecosystems with few hard barriers to dispersal that can facilitate allopatric speciation followed by secondary contact (e.g. marine habitats). Here we use a genomic (ddRADseq), model-based approach to delimit a cryptic species complex of tropical sea anemones that are co-distributed on coral reefs throughout the Tropical Western Atlantic. We use coalescent simulations in *fastsimcoal2* to test competing diversification scenarios that span the allopatric-sympatric continuum. We recover support that the corkscrew sea anemone *Bartholomea annulata* (Le Sueur, 1817) is a cryptic species complex, co-distributed throughout its range. Simulation and model selection analyses suggest these lineages arose in the face of historical and contemporary gene flow, supporting a sympatric origin, but an alternative secondary contact model also receives appreciable model support. Leveraging the genome of *Exaiptasia pallida* we identify five loci under divergent selection between cryptic *B. annulata* lineages that fall within mRNA transcripts or CDS regions. Our study provides a rare empirical, genomic example of sympatric speciation in a tropical anthozoan-a group that includes reef-building corals. Finally, these data represent the first range-wide molecular study of any tropical sea anemone, underscoring that anemone diversity is under described in the tropics, and highlighting the need for additional systematic studies into these ecologically and economically important species.

## 1. Introduction

Understanding the processes by which species form is fundamental to evolutionary biology. Historically, Darwin championed the role of deterministic mechanisms (i.e. selection), while the dominant paradigm of the Modern Synthesis required physical isolation as the primary starting point for reproductive isolation and speciation (e.g. Bird, Fernandez-Silva, Skillings, & Toonen, 2012; Bowen, Rocha, Toonen, Karl, & the ToBo Laboratory, 2013; Coyne & Orr, 2004; Gaither et al., 2015; Orr & Smith, 1998; Via, 2001). Under a deterministic framework, physical isolation is not a prerequisite for speciation, and selection can maintain reproductive isolation between sympatric populations in the early stages of divergence. The influential biologists of the 20^th^ century Modern Synthesis (i.e. Dobzhansky, Mayr, Maynard Smith) largely rejected sympatric speciation (reviewed by Bird et al., 2012), citing the homogenizing effects of gene flow and chromosomal rearrangement to prevent loci under divergent selection from accumulating and promoting reproductive isolation. Today, after decades of molecular data generation and increasing DNA sequencing technology, the idea of sympatric speciation is no longer as controversial. The gradually evolving genome championed by the Modern Synthesis is now giving way to a paradigm where it is understood that selection can drive divergence over short timescales without physical isolation, even in the face of historical and ongoing gene flow (e.g. Christie et al., 2017; Dennenmoser, Vamosi, Nolte, & Rogers, 2017; Feder, Egan, & Nosil, 2012; Nadeau et al., 2012; Renaut et al., 2013).

However, the relative contributions of allopatric and sympatric speciation in generating patterns of global biodiversity remain largely unresolved (e.g. Bolnick & Fitzpatric, 2007; Bowen et al., 2013) because different evolutionary processes can lead to similar outcomes. For example, it is challenging to empirically establish that co-distributed sister taxa diversified in sympatry versus allopatry, and that the contemporary geographic overlap isn’t the result of secondary contact following an allopatric diversification event (reviewed by Bird et al., 2012). Coyne and Orr (2004) suggest that divergence in sympatry, versus allopatric divergence and secondary contact, would be inferred only if the species exhibit reciprocal monophyly and are reproductively incompatible, or that an allopatric explanation seems unlikely. These criteria may be difficult to establish, as newly diverged species may not be reciprocally monophyletic across all individuals at all loci due to gene flow, periodic introgression, and incomplete lineage sorting. Further, contemporary distributions may not reflect historical ones, and range shifts in the distant past may obscure the geographic setting of diversification (e.g. Quenouille et al., 2011; Renema et al., 2008). Mathematical models present another paradigm to demonstrate that a sympatric explanation is favored over alternative scenarios (Bird et al., 2012). These allow researchers to test competing diversification scenarios, build models that incorporate explicit parameters for the directionality, magnitude, and timing of migration events, and use model selection to objectively and quantitatively inform demographic inference. For co-distributed sister taxa with no obvious geographic partitioning, patterns that demonstrate divergence in the face of ancestral and contemporary gene flow are considered among the strongest lines of evidence supporting sympatric diversification scenarios (Bird et al., 2012).

Tropical coral reefs are being increasingly viewed as fruitful ecosystems to explore sympatric speciation (Bowen et al., 2013). Coral reefs are the most biodiverse marine habitats on the planet, yet the bulk of diversity resides on less than 0.1% of the seafloor, in a setting with few hard barriers to dispersal. Purely allopatric models of speciation are an uneasy fit for describing diversification on this scale in this habitat (Bowen et al., 2013; Gaither et al., 2015; Gaither & Rocha, 2013; Rocha & Bowen, 2008). Evidence for non-allopatric divergence on coral reefs has been increasing, mainly from reef fishes, including species such as angelfishes, hamlets, damselfishes, wrasses, basslets, grunts, and gobies (Bernal, Gaither, Simison, & Rocha, 2017; Bowen et al., 2013; Hodge, Read, Bellwood, & Herwerden, 2013; Gaither et al., 2015; ; Munday, van Herwerden, & Dudgeon, 2004;). Examples from invertebrate species are rarer, but sympatric speciation has been invoked to explain to the diversification in limpets, nudibranchs, sponges, and corals (reviewed by Bowen et al. 2013). Most of the accepted examples of putative sympatric speciation from tropical marine systems have been made by documenting genetic differentiation between species/populations with overlapping distributions, but that segregate ecologically or in some other non-allopatric manner (e.g. Bongaerts et al., 2010; 2013). Other studies have used genomics to search for the basis of ecological adaptation for co-distributed species, and thus the basis for maintaining reproductive isolation and putative cause of divergence, but have not focused on the historical demographics of the speciation process itself (e.g. Rose, Bay, Morikawa, & Palumbi, 2018). The result is that many of the studies invoking sympatric diversification to explain the observed genetic and ecological patterns on reefs, while likely correct in many cases, arrive at these conclusions *post hoc*, and do not test competing speciation hypotheses. Modeling alternative processes provides a more objective way to assess the contributions of sympatric speciation, and helps avoid data over interpretation by incorporating statistical uncertainty into the model selection process (e.g. Knowles, 2009).

Here we conduct range-wide population-level sampling on coral reefs throughout the Tropical Western Atlantic (TWA) for the corkscrew sea anemone *Bartholomea annulata*, a common and ecologically important sea anemone that is described as a single species throughout its range (e.g. Briones-Fourzán, Pérez-Ortiz, Negrete-Soto, Barradas-Ortiz, & Lozano-Álvarez, 2012; Titus & Daly, 2017; Titus, Daly, & Exton, 2015; Titus et al., 2017). Using a double digest restriction-site associated DNA sequencing (ddRADseq) approach, we detect support for a previously unrecognized cryptic species that is co-distributed throughout the region. We then use the joint-folded allele frequency spectrum and coalescent simulations to model alternative diversification hypotheses. Finally, we conduct genome scans to identify loci under putative natural selection, and then leverage the close relationship between *B. annulata* and *Exaiptasia pallida* (see Grajales & Rodriguez, 2016), a species for which a genome has been published (Baumgarten et al., 2014), to explore whether these fall within, or are linked to, functional coding regions. Our model selection results provide one of the first genomic examples of sympatric speciation in the face of historical and ongoing gene flow in a tropical anthozoan, although an alternative secondary-contact model receives appreciable model support. This study highlights the importance of testing alternative diversification hypotheses and accounting for model uncertainty when conducting studies aimed at empirically demonstrating sympatric diversification. Lastly, these data represent the first range-wide molecular investigation into any reef-dwelling sea anemone in the world, underscoring that anemone diversity is under described in the tropics, and highlighting the need for additional systematic studies into these ecologically and economically important species.

## 2. Material and Methods

### 2.1. Sample collection, DNA isolation, and library preparation

Tissue samples (i.e. tentacle clippings and whole animals) were collected by hand using SCUBA from 14 sample localities spanning the geographic range of *B. annulata*, and from localities separated by known phylogeographic barriers (Fig. 1; reviewed by DeBiasse, Richards, Shivji, & Hellberg, 2016). Samples were collected from coral reef habitats between 5-and 20-m depth, preserved on shore, and transferred back to The Ohio State University for DNA extraction and sequencing. Genomic DNA was isolated using DNeasy Blood and Tissue Kits (Qiagen Inc.) and stored at −20°C. DNA degradation was assessed for each sample using gel electrophoresis, and only samples with high molecular weight DNA were carried forward for ddRADseq library preparation. DNA concentrations were quantified (ng/uL) using a Qubit 2.0 (ThermoFisher) fluorometer and dsDNA broad-range assay kits. 20uL aliquots, each with 200ng of DNA, were prepared for each sample and used for ddRADseq library preparation.

**Figure 1.**
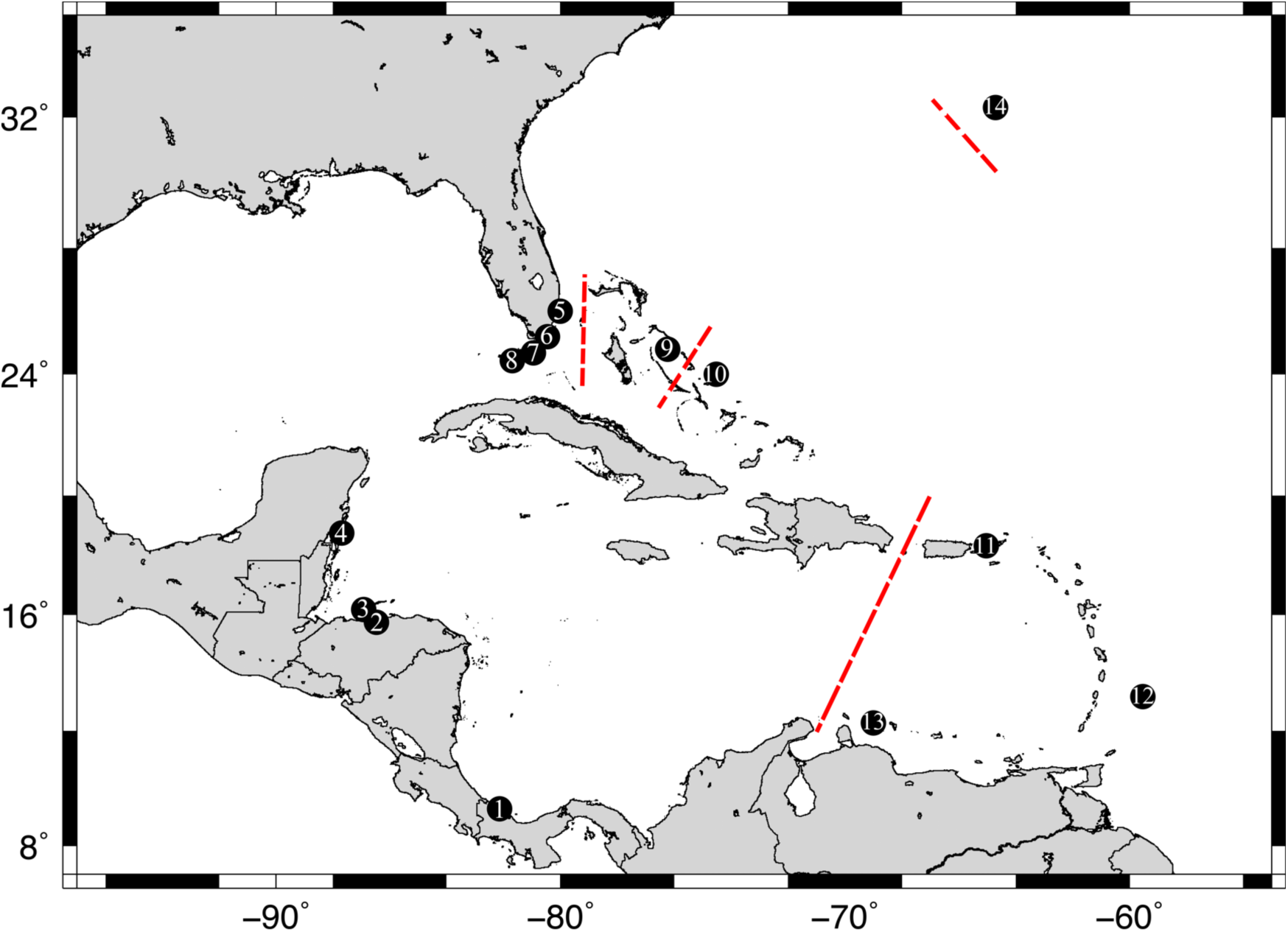
Map of sampling localities in the Tropical Western Atlantic for the corkscrew sea anemone *Bartholomea annulata*. 1) Bocas del Toro, Panama, 2) Cayos Cochinos, Honduras, 3) Utila, Honduras, 4) Mahahual, Mexico, 5) Ft. Lauderdale, Florida, 6) Upper Keys, Florida, 7) Middle Keys, Florida, 8) Lower Keys, Florida, 9) Eleuthera, Bahamas, 10) San Salvador, Bahamas, 11) St. Thomas, US Virgin Islands, 12) Barbados, 13) Curacao, 14) Bermuda. Dashed lines denote previously recovered major phylogeographic breaks and allopatric boundaries in the region. Sympatric lineages were recovered from all localities except Honduras (2, 3), Virgin Islands (11), and Bermuda (14).

Between 12-15 individual *B. annulata* samples per locality were carried forward for library preparation. We used a ddRADseq library preparation protocol following Sovic, Fries, & Gibbs (2016). Briefly, genomes were digested using two restriction enzymes (*Eco*RI-HF and *psti*-HF), Illumina compatible barcodes were annealed to restriction cut sites, samples were size selected manually using a 400-800 bp size range, and then cleaned using Nucleospin Gel and PCR clean up kits (Macherey-Nagel). Following size selection, each individual sample was amplified using polymerase chain reaction (PCR), cleaned using AMpure XP beads (Agilent), and then quantified via quantitative PCRs (qPCR) to inform the pooling of individual samples into final libraries. A total of 141 individuals (Table 1) met all quality control steps and were pooled across five separate libraries. Samples were sequenced on an Illumina HiSeq 2500 using single-end 100 base pair reads at The Ohio State University Genomics Shared Resource.

**Table 1.** Sample localities, sample sizes, and geographic coordinates of corkscrew sea anemone *Bartholomea annulata* used in this study. Sample sizes reflect the number samples sequenced and the number of samples retained in the final double digest Restriction-site Associated DNA sequencing (ddRADseq) dataset (in parentheses). Differences between the number of samples sequenced and retained reflects variation in the number of sequence reads and sequencing coverage in out ddRADseq dataset across all individuals.

| Locality | Code | Sample size | Latitude | Longitude |
| --- | --- | --- | --- | --- |
| Eleuthera, Bahamas | BH | 10 (9) | 24°49'44.51"N | 76°16'46.11"W |
| San Salvador, Bahamas | SAN | 10 (2) | 24° 2'37.12"N | 74°31'59.83"W |
| Barbados | BR | 10 (9) | 13°11'30.52"N | 59°38'29.04"W |
| Bermuda | BD | 12 (9) | 32°26'53.62"N | 64°45'45.42"W |
| Curacao | CU | 10 (2) | 12° 7'19.45"N | 68°58'10.80"W |
| Ft. Lauderdale, Florida, USA | FT | 9 (9) | 26° 4'19.80"N | 80° 5'46.68"W |
| Upper Keys, Florida, USA | UK | 10 (9) | 25° 1'57.92"N | 80°22'4.45"W |
| Middle Keys, Florida, USA | MK | 10 (10) | 24°41'58.09"N | 80°56'21.48"W |
| Lower Keys, Florida, USA | LK | 10 (10) | 24°33'42.39"N | 81°23'31.59"W |
| Utila, Honduras | UT | 10 (10) | 16° 5'18.03"N | 86°54'38.54"W |
| Cayos Cochinos, Honduras | CC | 10 (9) | 15°57'1.12"N | 86°29'51.82"W |
| Mahahual, Mexico | MX | 10 (10) | 18°42'18.45"N | 87°42'34.46"W |
| Bocas del Toro, Panama | PA | 10 (10) | 9°25'7.28"N | 82°20'32.55"W |
| St. Thomas, US Virgin Islands | VI | 11 (11) | 18°19'0.69"N | 64°59'22.59"W |

### 2.2. Data processing and dataset assembly

Raw sequence reads were demultiplexed, aligned, and assembled *de novo* using the program pyRAD v3.0.66 (Eaton, 2014). We required a base call Phred score of 20 and set the maximum number of bases in a locus with Phred scores < 20 (NQual) to five. Low quality base calls were replaced with Ns. We set the clustering threshold (Wclust) to 0.90 to assemble reads into loci, and required a minimum coverage depth of seven to call a locus (Mindepth). Finally, we required a locus to be present in 75% of all individuals to be retained in the final dataset. RADseq protocols are known to be susceptible to missing data due to mutations in restriction cut sites and allelic dropout (e.g. Arnold, Corbett-Detig, Hartl, & Bomblies, 2013), but biases can also arise when datasets are overly conservative (i.e. no missing data allowed; Huang & Knowles, 2014). Thus we allowed some missing data in our final dataset.

Like most coral reef dwelling anthozoans, *B. annulata* hosts endosymbiotic dinoflagellates from the genus *Symbiodinium* (e.g. Grajales & Rodriguez, 2016). Thus, DNA extractions harbor a mix of anemone and dinoflagellate DNA, and the resulting ddRAD sequencing yields a mixture of anemone and *Symbiodinium* sequences. To deal with the potentially confounding *Symbiodinium* DNA contamination and verify that our final data set contains SNPs that are anemone DNA only, we created a symbiont-free dataset by mapping ddRADseq loci to the *Exaiptasia pallida* genome (Baumgarten et al., 2014). *Exaiptasia pallida* and *B. annulata* are members of the same family and are closely related (Grajales & Rodriguez, 2016), and polymorphic microsatellites have previously been designed from *E. pallida* that amplify in *B. annulata* (Titus et al., 2017). To map polymorphic *B. annulata* loci to *E. pallida*, we downloaded the *E. pallida* genome and created a local BLAST database. After initially running pyRAD to completion, a python script (parse_loci.py, available on Dryad doi:XXX) was written to select the first DNA sequence from each locus in the .loci output file, and create a .fasta file that could then be BLASTed against the *E. pallida* genome (BLAST+; Camacho et al, 2009). We used an 85% identity threshold to call a locus as putatively anemone in origin, to avoid being over conservative as we were not mapping to a conspecific genome. Next, a separate python script (blast2loci.py, available on Dryad doi:XXX) was written to read through the BLAST output file, pull all sequences in all loci that met the 85% identity threshold, and create a new .loci file with the same file name as the original. The original .loci file was then replaced with the new anemone-only file, at which point the final step of pyRAD (step 7) was re-run to create our final anemone-only output files (i.e. unlinked SNPs and alleles files) for downstream analyses.

### 2.3. Genetic clustering and species delimitation

To search for evidence of cryptic species-level diversity, we used the clustering program Structure v2.3.4 (Pritchard, 2000) as a preliminary species discovery analysis (i.e. no *a priori* species assignments; reviewed by Carstens, Pelletier, Reid, & Satler, 2013). We collapsed bi-allelic data into haplotypes at each locus, using information contained in linked SNPs when more than one SNP was present in a locus. Each analysis in Structure was run using the full set of samples (Table S1), and used the admixture model, correlated allele frequencies, and sampling location. Each MCMC chain for each value of *K* was run with a burnin of 1 x 10^5^ generations and sampling period of 2 x 10^5^ generations. We conducted five iterations of a broad range of *K* values (1–6), to gain an initial snapshot of the data across the region. In both initial analyses we used the peak ln Pr(*D*|*K*) and the Δ*K* (Evanno et al. 2005) to help select the best *K* value.

Structure analyses overwhelmingly selected *K* = 2 as the best clustering scheme with both genetic clusters being co-distributed throughout the entire TWA, save for Bermuda and the US Virgin Islands (see Results). Because this pattern may suggest the presence of an unrecognized cryptic anemone species, we conducted species delimitation analyses using path sampling and Bayes factors using the program SNAPP and Bayes Factor Delimitation* (BFD*; Leache, Fujita, Minin, & Bouckaert, 2014). BFD* uses unlinked SNPs and marginal likelihood estimates (MLEs) calculated via path sampling in the species tree program SNAPP (Bryant et al., 2012) to perform model selection on competing species delimitation models. We tested the current concept of *B. annulata* (i.e. a single TWA species) versus the alternative model (i.e. two sympatric species). Bayes Factors were calculated after Leache et al. (2014). Positive values indicate support for model 1 (current concept) while negative values indicate support for model 2 (competing species delimitation models; Leache et al., 2014).

Due to the computational constraints running SNAPP with biallelic SNP data (i.e. computation time increases linearly as more loci are added but exponentially as more samples are added), and missing data constraints for estimating species trees (i.e. data has to be present in at least one individual at each locus for each putative species) we created new SNP datasets by significantly reducing the number of total individuals so that our analyses could be completed over the course of days rather than weeks or months. Similar approaches have been taken by previous studies (e.g. Sovic et al., 2016). We used n = 5 randomly selected individuals from each genetic cluster delimited by Structure and used *E. pallida* as an outgroup as BFD* can only perform model selection on models with n ≥ 2 species (Table S2).

Of 11 total individuals included in this subset analysis, we required a locus to be present in 10 of 11 individuals to meet the requirements of species tree estimation. Our species delimitations were thus a two species model (*E. pallida* + current concept of *B. annulata*) and a three species model (*E. pallida* + *B. annulata* Clade 1 + *B. annulata* Clade 2). For each SNAPP analysis, mutation rates *u* and *v* were set to 1 and were not sampled and the coalescent rate was set at 10 and sampled throughout the analysis. We used only polymorphic loci and a broad gamma distributed (2, 200) prior for speciation rate (*λ*). Each step in the path analysis (48 steps) was conducted in SNAPP v1.3 and BEAST v.2.4.6 (Bouckaert et al., 2014) for 1 x 10^5^ MCMC generations with 10% discarded as burnin.

As we did not include any *E. pallida* samples in our ddRAD sequencing, we wrote a python script that located and pulled the segments of DNA from the *E. pallida* genome where our 100bp *B. annulata* loci mapped with high confidence using BLAST searches (add_outgroup.py, available on Dryad; LINK). These 100bp segments of *E. pallida* DNA were then incorporated into, and aligned with, our *B. annulata* RAD loci using Muscle v3.8.31 (Edgar 2004). SNPs were recoded and one SNP per locus was pulled randomly to create a new unlinked SNP .nexus file that could be inputted into SNAPP (aln2snapp.py, available on Dryad doi:XXX). This novel approach for using current genomic resources to add outgroups to RADseq datasets should be amenable to any set of closely related species. The script, along with full details and instructions for using it, can be found on Dryad (doi:XXX).

### 2.4. Model selection

We used the allele frequency spectrum (AFS) and a coalescent simulation approach using the program *fastsimcoal2* (FSC2; Excoffier, Dupanloup, Huerta-Sánchez, Sousa, & Foll, 2013) to provide empirical support that the pattern of cryptic species-level diversity we detect using our ddRADseq dataset most likely arose sympatrically. FSC2 uses coalescent simulations to calculate the composite likelihood of arbitrarily complex demographic models under a given AFS. The best-fit model can then be selected using the Akaike Information Criterion (AIC).

We built 12 demographic models (Fig. 2), all variants of the two-population isolation-migration models, as Structure and BFD* analyses support two lineages of *B. annulata* (see Results). Models are isolation-only, isolation followed by secondary contact, and models that incorporate historical and contemporary gene flow (Fig. 2). We aim to test the likelihood of alternative hypotheses given the available data rather than attempting to prove sympatric speciation with “air-tight” evidence (Bird et al., 2012). For lineages that are co-distributed throughout their entire range, the strongest evidence for sympatric diversification would be models that demonstrate both historical and contemporary gene flow (i.e. no interruption of gene flow). These models necessarily exclude allopatric scenarios where complete geographic isolation has disrupted gene flow to initiate divergence (Bird et al., 2012). Conversely, the weakest evidence for sympatric speciation would be isolation-only and secondary-contact models. In the latter scenario, we would fail to reject scenarios where divergence occurred with complete allopatric isolation but became sympatrically distributed and resumed gene flow after secondary contact.

**Figure 2.**
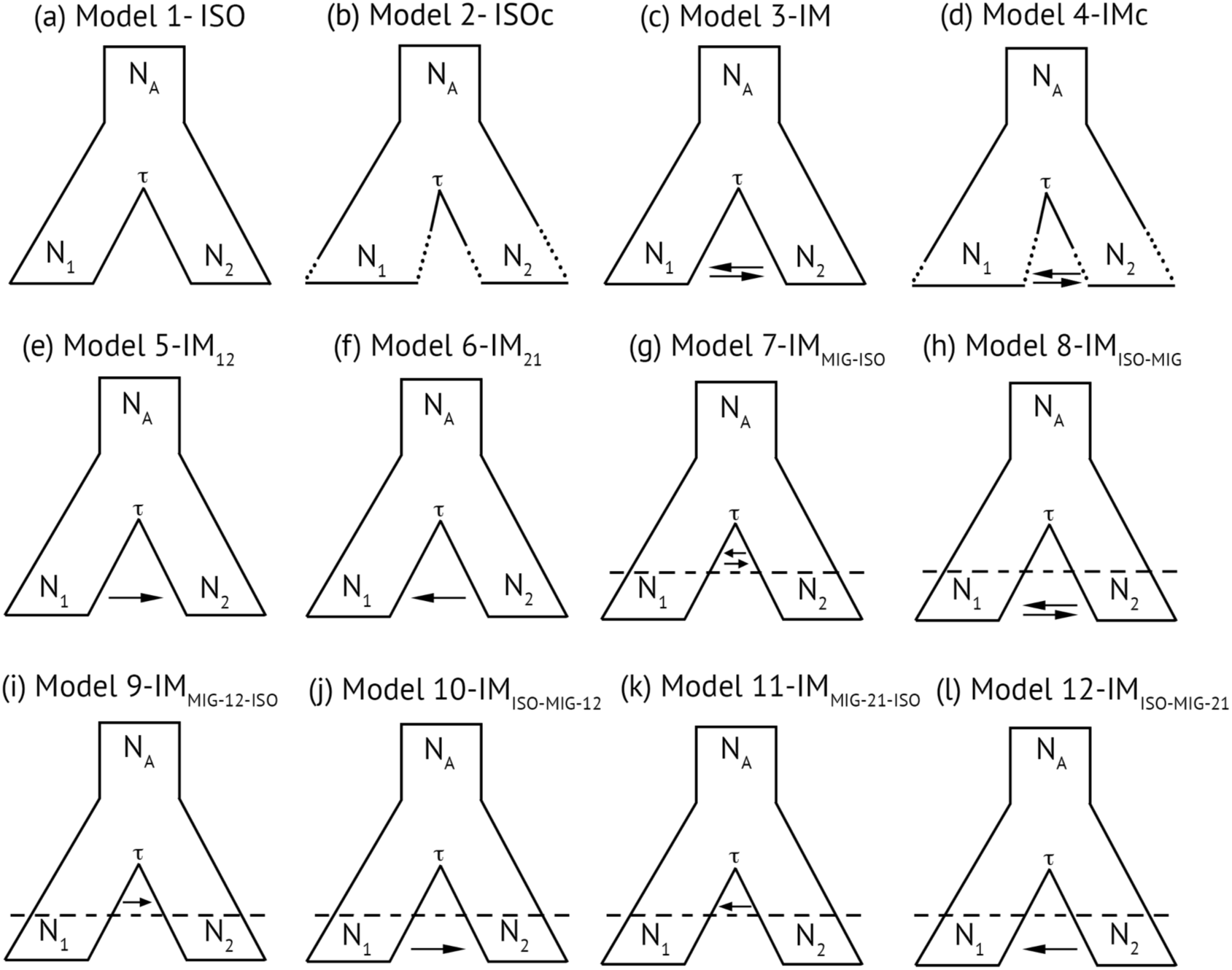
Models used in FSC2 to understand the demographic processes leading to cryptic diversification in the corkscrew anemone *Bartholomea annulata*. Each model is a two-population isolation-migration (IM) model that varies in the degree and directionality of gene flow and effective population size. Models are as follows: a) isolation only, b) isolation only with population size changes following divergence, c) IM model with symmetric migration, d) IM model with symmetric migration and population size changes, e) IM model with migration from population 1 to 2, f) IM model with migration from population 2 to 1, g) IM model with symmetric migration between populations immediately following divergence, followed by more recent isolation, h) IM model with isolation immediately following divergence, followed by more recent secondary contact and symmetric migration, i) IM model with migration from population 1 to 2 immediately following divergence followed by more recent isolation, j) IM model with isolation immediately following divergence followed by secondary contact and migration from population 1 to 2, k) IM model with migration from population 2 to 1 immediately following divergence followed by more recent isolation, and l) IM model with isolation immediately following divergence followed by secondary contact and migration from population 2 to 1.

To conduct simulation analyses, two-population, joint-folded AFS were generated from pyRAD output files and previously published python scripts (see Satler & Carstens, 2017) using 24 randomly selected individuals from the more well sampled *B. annulata* lineage (Clade 2) and all 16 individuals from the less sampled lineage (Clade 1; Table S3). One of the assumptions of FSC2 is that SNPs are in linkage equilibrium (Excoffier et al., 2013), and thus, only one SNP per locus was selected to produce the AFS. Further, AFS calculations in FSC2 require fixed numbers of alleles from all populations (i.e. no missing data). As meeting this latter requirement would greatly decrease our dataset size, and thus likely bias our analyses, we followed the protocol of Satler and Carstens (2017) and Smith et al., (2017) by requiring a locus in our AFS to be present in 85% of all individuals. To account for missing data without violating the requirements of the AFS we built our AFS as follows: 1) if a locus had fewer alleles than our threshold it was discarded, 2) if a locus had the exact number of alleles as the threshold, the minor allele frequency was recorded, and 3) if a locus exceeded the threshold, alleles were down-sampled with replacement until the number of alleles met the threshold, at which point the minor allele frequency was counted. This approach allowed us to maximize the number of SNPs used to build the AFS, but also has the potential to lead to monomorphic alleles based on the down-sampling procedure (see Satler and Carstens 2017). Thus, we repeated the AFS building procedure 10 times, allowing us to account for variation in the down-sampling process during model selection, but also allowing us to calculate confidence intervals on our parameter estimates (Satler and Carstens, 2017; Smith et al., 2017.

Each simulation analysis in FSC2 (i.e. each AFS replicate per model; 12 models x 10 replicates) was repeated 50 times, and we selected the run with the highest composite likelihood for each AFS replicate and model. The best-fit model was then calculated using the AIC and model probabilities calculated following Burnham and Anderson (2002). Because FSC2 requires a per generation mutation rate to scale parameter estimates into real values, we used the substitution per site per generation mutation rate of 4.38 x 10^-8^proposed for tropical anthozoans (Prada et al., 2017) and a generation time of 1 year for *B. annulata* (Jennison, 1981). All analyses were conducted on the Oakley cluster at the Ohio Supercomputer Center (http://osc.edu).

### 2.5. Detecting and identifying loci under selection

Islands of genomic divergence linked to functional genes under selection are common in sympatrically diverging species (e.g. Renaut et al., 2013). To detect and identify loci under putative selection that may be contributing to the divergence of our newly delimited *B. annulata* lineages we used the program BayeScan v.2.1 (Foll & Gaggiotti, 2008). This program uses logistical regression and a Bayesian framework to statistically search for loci under natural selection. It implements a locus effect and population effect in the model to explain the observed patterns of differentiation. If the locus effect is needed to explain the pattern, divergent selection is indicated (Folk & Gaggiotti, 2008). We used 100,000 simulations with prior odds of 8 (i.e. how much more likely the neutral model is than the selection model), and a false discovery rate of 10%. Results were summarized and outlier loci visualized using R v3.3.1 and RStudio v0.98 (R Core Team 2014).

We used the genome of *E. pallida* to identify where in the genome the SNPs under putative selection reside and to determine whether they are in, or in close proximity to, functional coding regions. We used the full 100bp sequence from which each outlier SNP was recovered, and then mapped each locus to the annotated *E. pallida* genome (Baumgarten et al., 2014) using Geneious v10.2.3 (Kearse et al., 2012). For outlier loci that did not fall within coding regions, we recorded the distance (in bp) to the next-closest coding region, as these loci may be linked to functional genes under selection.

## 3. RESULTS

### 3.1. RADseq dataset assembly

Double digest RADseq library preparation and sequencing resulted in a total of 186.7 million sequence reads across 141 individuals, 175.6 million of which passed quality control filtering and were retained to create the final dataset. Twenty-two of the 141 individuals had < 500,000 reads and were not retained in the final dataset. Requiring a locus to be present in a minimum of 75% of all individuals resulted in a final data set of 10,998 parsimoniously informative sites distributed across 3176 unlinked loci in 119 individuals. BLASTing these loci to the *Exaiptasia pallida* genome identified 1402 loci that matched with high confidence (≥ 85% identity); these were used as the final anemone-only SNP dataset. The remaining 1772 loci that did not map to any genomic resources were discarded as their identity (anemone or algal) could not be verified. SNP files and datasets are available on Dryad (Dryad doi:XXX).

### 3.2. Genetic clustering and species delimitation

Genetic clustering approaches in Structure [i.e. Δ*K* and the mean lnP(*K*)] both selected *K* = 2 as the optimum partitioning scheme (Table 2; Figure 3). Both genetic clusters were co-distributed and recovered from all sampling localities with the exception of Bermuda, Honduras, and the US Virgin Islands. One cluster, henceforth *B. annulata* Clade 1, was sampled infrequently and was represented by only 16 individuals (13% of all sampled individuals) throughout the TWA, while the second cluster, henceforth *B. annulata* Clade 2, was well sampled and comprised the majority of the samples (87%; Figure 3). Some admixture between *B. annulata* clades is evident in our Structure results (Figure 3). Species delimitation analyses using SNAPP and BFD* support both genetic clusters recovered by Structure as separate species, favoring the alternative model to the current taxonomy model (Table 3).

**Table 2.** Results of Structure analyses for the corkscrew anemone *Bartholomea annulata*. *K* is the number of genetic clusters tested for each model. Highlighting indicates the model with the best support, as determined by the mean natural log posterior probabilities [lnP(*K*)] and delta *K* (Δ*K*).

| $K$ | Iterations | mean $\ln P(K)$ | Stdev $\ln P(K)$ | $\Delta K$ |
| --- | --- | --- | --- | --- |
| 1 | 5 | -535786.16 | 18.47 | N/A |
| 2 | 5 | -194732.23 | 27.09 | 12637.83 |
| 3 | 5 | -196054.70 | 249.66 | 4.42 |
| 4 | 5 | -196272.33 | 738.44 | 0.99 |
| 5 | 5 | -197221.50 | 2098.35 | 36.97 |
| 6 | 5 | -275748.30 | 135217.75 | N/A |

**Table 3.**
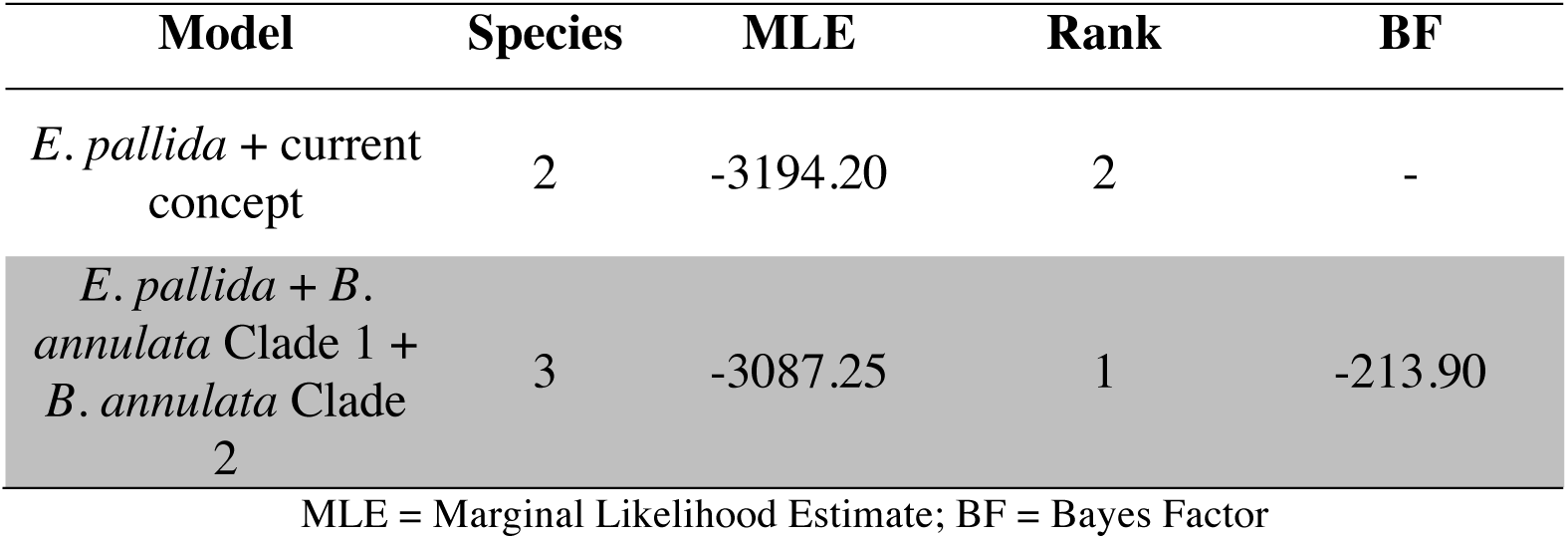
Path sampling results for two species delimitation models for *Bartholomea annulata* in the Tropical Western Atlantic. All Bayes Factor calculations are made against the current model describing *B. annulata* as a single species. Positive Bayes Factors indicate support for the current model. Negative Bayes Factors indicate support for the alternative model, in which *B. annulata* comprises two species. For both models, *Exaiptasia pallida* was included as an outgroup. The higher ranked model is highlighted.

**Figure 3.**
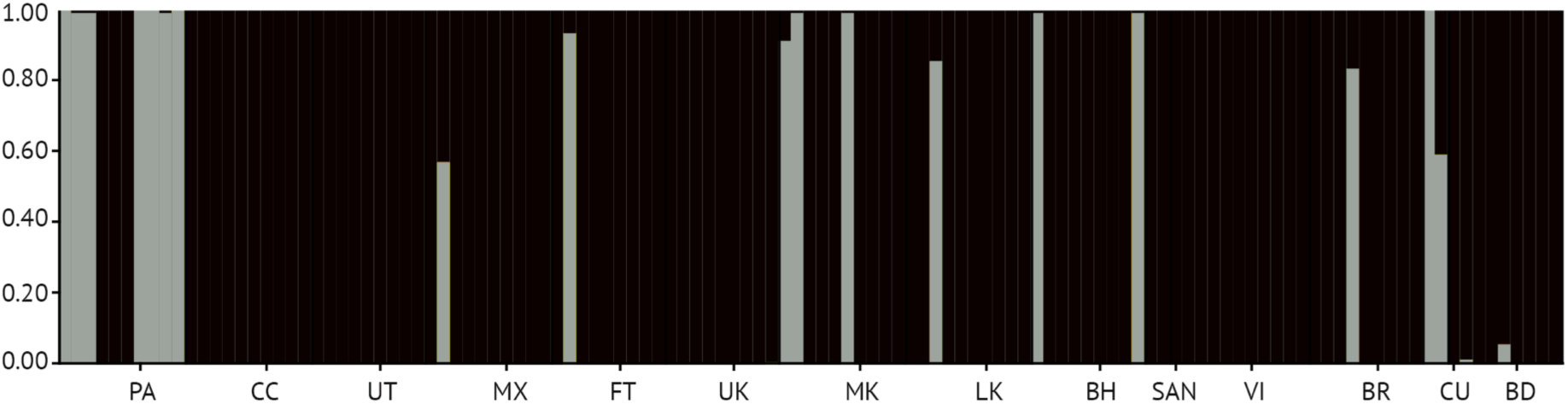
Structure plot denoting *K* = 2 genetic clusters for the corkscrew sea anemone *Bartholomea annulata* distributed sympatrically throughout the Tropical Western Atlantic. Samples are partitioned by sample locality (Fig 30): PA = Bocas del Toro, Panama, CC = Cayos Cochinos, Honduras, UT = Utila, Honduras, MX = Mahahual, Mexico, FT = Ft. Lauderdale, Florida, UK = Upper Keys, Florida, MK = Middle Keys, Florida, LK = Lower Keys, Florida, BH = Eleuthera, Bahamas, SAN = San Salvador, Bahamas, VI = St. Thomas, US Virgin Islands, BR = Barbados, CU = Curacao, BD = Bermuda. The order of listed sample localities reflects a roughly West to East distribution across the Tropical Western Atlantic.

### 3.3. Coalescent model selection

The best fit model under the Akaike Information Criterion (AIC) is model 5 (Figure 2), an IM model with unidirectional gene flow from *B. annulata* Clade 1 to Clade 2 (Table 4). Also supported was model 10, an IM model with isolation immediately following divergence followed by secondary contact and migration from *B. annulata* Clade 1 to Clade 2. Each of the top three models according to AIC specified unidirectional gene flow from *B. annulata* Clade 1 to Clade 2. Isolation-only models received an inconsequential amount of support (Table 4). For the best-fit model, FSC2 simulation places divergence time estimates between putative species at 2.1 mya. In general, parameter estimates for the two best-fit models were broadly similar, except in divergence time, which was estimated at a far older date in model 10 than model 5 (Table 5).

**Table 4.** Akaike Information Criterion (AIC) results for model selection results from FSC2. Model refers to those depicted and described in Figure 2. *k* = number of parameters in the model, Δ_I_ = change in AIC scores, and *w*_*i*_ = Akaike weights. Models are listed according to their AIC rank and the highest-ranked model is highlighted.

| Model | $k$ | $\ln(\text{Likelihood})$ | AIC | $\Delta_i$ | Model Likelihoods | $w_i$ |
| --- | --- | --- | --- | --- | --- | --- |
| 5 – IM<sub>12</sub> | 5 | -2285.63 | 4581.27 | 0 | 1 | 0.65 |
| 10 – IM<sub>ISO-MIG-12</sub> | 6 | -2285.52 | 4583.04 | 1.76 | 0.41 | 0.27 |
| 9 – IM<sub>MIG-12-ISO</sub> | 6 | -2287.03 | 4586.07 | 4.79 | 0.09 | 0.06 |
| 7 – IM<sub>MIG-ISO</sub> | 7 | -2287.24 | 4588.49 | 7.21 | 0.02 | 0.02 |
| 8 – IM<sub>ISO-MIG</sub> | 7 | -2293.17 | 4600.35 | 19.07 | 7.20e<sup>-5</sup> | 4.70e<sup>-5</sup> |
| 3 – IM | 6 | -2294.36 | 4600.72 | 19.44 | 5.97e<sup>-5</sup> | 3.90e<sup>-5</sup> |
| 1 – ISO | 4 | -2309.26 | 4626.52 | 45.25 | 1.49e<sup>-10</sup> | 9.73e<sup>-11</sup> |
| 6 – IM<sub>21</sub> | 5 | -2322.01 | 4654.03 | 72.75 | 1.58e<sup>-16</sup> | 1.03e<sup>-16</sup> |
| 11 – IM<sub>MIG-21-ISO</sub> | 6 | -2357.18 | 4726.37 | 145.09 | 3.11e<sup>-32</sup> | 2.03e<sup>-32</sup> |
| 12 – IM<sub>ISO-MIG-12</sub> | 6 | -2362.40 | 4736.80 | 155.52 | 1.69e<sup>-34</sup> | 1.10e<sup>-34</sup> |
| 2 – ISC<sub>C</sub> | 11 | -2436.15 | 4894.31 | 313.03 | 1.06e<sup>-68</sup> | 6.92e<sup>-69</sup> |
| 4 – IM<sub>C</sub> | 12 | -2557.29 | 5138.59 | 557.31 | 9.55e<sup>-122</sup> | 6.23e<sup>-122</sup> |

**Table 5.** Population genetic parameter estimates from FSC2 for the two cryptic lineages of *Bartholomea annulata* (Clade 1 & Clade 2) for the two demographic models with the highest Akaike model weights from Table 4. Parameter estimates generated with unlinked allele frequency spectrum data. Divergence time (*τ*) and time at secondary contact (*τ*-*M*) are in millions of years (mya), effective population sizes (N_*e*_) are presented as the effective number of individuals in each population, and migration rates (*M*) are presented as the effective number of migrants per generation. N/A refers to parameters that were not included in the model.

| **Model** | **$\tau$ (mya)** |  | **$N_e$ Clade1** |  | **$N_e$ Clade2** |  | **$\tau-M$** |  | **$M_{12}$** |  |
| --- | --- | --- | --- | --- | --- | --- | --- | --- | --- | --- |
|  | Mean | 95% CI | Mean | 95% CI | Mean | 95% CI | Mean | 95% CI | Mean | 95% CI |
| **5** | 2.11 | ($\pm 1.62$) | 23.70 | ($\pm 10.4$) | 107,348 | ($\pm 5339$) | N/A | N/A | $4.7e^{-5}$ | ($\pm 6.04 e^{-6}$) |
| **10** | 8.87 | ($\pm 3.20$) | 45.40 | ($\pm 5302$) | 98,213 | ($\pm 10,619$) | 2.10 | ($\pm 2.05$) | $5.5e^{-5}$ | ($\pm 1.63 e^{-5}$) |

### 3.4. Detecting and identifying loci under selection

We detected 31 outlier loci, out of 1401 total loci, using BayeScan v.2.1 (Figure S1). Six were characterized as being outliers with high Fst, and putatively under divergent selection. The remainder had low Fst, compared with the remaining dataset, and are putatively under balancing selection. The genomic positions of all 31 outlier loci were identified by mapping full 100 bp reads to the *E. pallida* genome. 15 loci mapped within exons of coding regions (CDS), nine mapped to mRNA transcripts, and the remaining loci did not map to any known functional regions (Table 6). Of the loci that did not map to an mRNA transcript or coding region, we identified the next closest gene and mRNA transcript, as these could potentially be linked to genes under selection.

**Table 6.** List of *Bartholomea annulata* outlier loci detected by BayeScan, the genome scaffold number they map to in the *Exaiptasia pallida* genome, whether they fall within an annotated region of the genome, and their closest functional gene identity. Loci identified as being within coding regions (CDS) and exons map <u>within</u> the specific gene listed. Loci identified as mRNA transcripts are listed beside the functional gene that corresponds with that transcript. Loci listed as N/A do not map to any known functional regions. Instead, the closest functional gene in the *E. pallida* genome is listed to the right. Highlighted loci (1, 97, 197, 1160, 1224, and 1326) are those under divergent selection (high Fst).

| Locus | Genome scaffold # | Annotation | Closest functional gene |
| --- | --- | --- | --- |
| 1 | 54 | mRNA | *E. pallida*: neuronal acetylcholine receptor subunit alpha-10-like-PREDICTED |
| 10 | 83 | mRNA/3' UTR | *E. pallida*: peroxisomal coenzyme A diphosphatase NUDT7-like-PREDICTED |
| 19 | 237 | N/A | *E. pallida*: inactive rhomboid protein 1-like-PREDICTED |
| 27 | 28 | mRNA/5' UTR | *E. pallida*: ubiquitin carboxyl-terminal hydrolase 8-like-PREDICTED |
| 97 | 14 | mRNA/3' UTR | *E. pallida*: C-C chemokine receptor type 4-like-PREDICTED |
| 142 | 86 | CDS/exon | *E. pallida*: twisted gastrulation protein homolog-PREDICTED |
| 197 | 197 | CDS/exon | *E. pallida*: zinc finger FYVE domain-containing protein 1-like-PREDICTED |
| 265 | 100 | CDS/exon | *E. pallida*: dual serine/threonine & tyrosine protein kinase-like-PREDICTED |
| 310 | 407 | CDS/exon | *E. pallida*: melatonin receptor type 1A-like-PREDICTED |
| 321 | 25 | CDS/exon | *E. pallida*: protocadherin-like protein-PREDICTED |
| 399 | 66 | CDS/exon | *E. pallida*: posphatidylineositide phosphate SAC2-like-PREDICTED |
| 522 | 233 | CDS/exon | *E. pallida*: uncharacterized protein |
| 602 | 358 | N/A | *E. pallida*: fibropellin-1-like transcript variant X1-PREDICTED |
| 684 | 138 | CDS/exon | *E. pallida*: YTH domain containing family protein 1-like-PREDICTED |
| 725 | 64 | mRNA/5'UTR | *E. pallida*: uncharacterized protein |
| 744 | 62 | CDS/exon | *E. pallida*: NAD-dependent progein deacetylase SRT1-like-PREDICTED |
| 757 | 78 | mRNA | *E. pallida*: dnaJ homolog subfamily C member 11-like-PREDICTED |
| 766 | 197 | N/A | *E. pallida*: zinc metalloproteinase nas-13-like-PREDICTED |
| 776 | 2486 | N/A | *E. pallida*: formin binding protein 4-like-PREDICTED |
| 814 | 542 | N/A | *E. pallida*: zinc finger protein 567-like-PREDICTED |
| 851 | 764 | mRNA | *E. pallida*: uncharacterized protein |
| 1013 | 99 | CDS/exon | *E. pallida*: pre-mRNA-processing-splicing factor 8-PREDICTED |
| 1017 | 1177 | mRNA | *E. pallida*: Golgi-associated plant pathogenesis-related protein 1-like-PREDICTED |
| 1043 | 155 | CDS/exon | *E. pallida*: GDAP2 protein homolog-PREDICTED |
| 1131 | 109 | N/A | *E. pallida*: uncharacterized |
| 1159 | 211 | CDS/exon | *E. pallida*: germinal center kinase 3-like-PREDICTED |
| 1160 | 210 | CDS/exon | *E. pallida*: snaclec rhodocetin subunit alpha-like-PREDICTED |
| 1213 | 5 | CDS/exon | *E. pallida*: uncharacterized protein |
| 1224 | 210 | N/A | *E. pallida*: receptor-type tyrosine-protein phosphatase delta-like-PREDICTED |
| 1312 | 19 | CDS/exon | *E. pallida*: nesprin-1-like-PREDICTED |
| 1326 | 30 | mRNA | *E. pallida*: phosphatase 1 regulatory subunit 12A-like-PREDICTED |

The six loci identified as being under divergent selection (loci 1, 97, 197, 1160, 1224, and 1326) all had Fst values exceeding 0.60 with a maximum observed Fst value of 0.93 (Figure S1). Locus 197 and 1160 were identified as residing within CDS/exon regions, and mapped to the predicted functional genes zinc finger FYVE domain-containing protein and snaclec rhodocetin subunit alpha respectively (Table 6). Locus 1, 97, and 1326 mapped to mRNA transcripts in association with neuronal acetylcholine receptor subunit alpha-10, C-C chemokine receptor type 4, and phosphate 1 regulatory subunit 12A genes respectively (Table 6). Locus 1224 did not map to a known functional region in the *E. pallida* genome and was 7,000 bp from a receptor-type tyrosine-protein phosphate delta gene.

## 4. DISCUSSION

### 4.1. Genomic signatures of sympatric speciation

Tropical coral reefs have extraordinary levels of biodiversity that reside on a small fraction of suitable habitat space and that occupy large biogeographic ranges. Most reef species have planktonic larval stages, and all marine species reside in a physical environment that should facilitate dispersal, gene flow, and large effective population sizes. This dichotomy highlights the challenge of disentangling sympatric diversification from allopatric diversification followed by secondary contact on reefs, as incipient species that diverge in allopatry can readily become co-distributed with their sister taxa unless permanently isolated by hard barriers to dispersal (i.e. continental land masses). Here we provide one of the first genomic, model-based, tests of sympatric speciation in a reef-dwelling species. Our coalescent simulation analyses and model selection suggest that two co-distributed lineages of sea anemones, currently described as a single species, the corkscrew anemone *Bartholomea annulata*, have diverged in the face of continuous, unidirectional, gene flow on reefs throughout the Tropical Western Atlantic. We note, however, that there is model support for an alternative secondary contact scenario where gene flow was suspended for a prolonged period of time.

Definitions of sympatric speciation have, historically, been contentious, and there is no agreed upon consensus definition (reviewed by Bird et al., 2012). We define sympatric speciation in a broad biogeographic sense: both lineages of *B. annulata* are sympatric in their broad-scale biogeography, co-occur in several places, and lack major allopatric barriers across their ranges. Well-resolved allopatric boundaries in the Tropical Western Atlantic linked to putative speciation events include the Mona Passage between the islands of Hispanola and Puerto Rico, the Florida Straits separating the Florida peninsula from the Bahamas and Cuba, the isolated archipelago of Bermuda, and a Central Bahamas break (reviewed by DeBiasse et al., 2016). Reefs off the coast of Panama are another biogeographic sub-region within the Tropical Western Atlantic that are physically isolated by ocean currents (i.e. the Panamanian Gyre; Richardson, 2005) and have a propensity to show limited genetic connectivity with nearby reefs (e.g. Andras, Kirk, & Harvell, 2011; Andras, Rypien, & Harvell, 2013; DeBiasse et al., 2016). Interestingly, 7 of the 16 individuals from our infrequently sampled Clade 1 lineage came from reefs from Bocas del Toro, Panama (Fig. 3). Given the appreciable model support for a secondary contact divergence scenario, it could be possible that a population of *B. annulata* became physically isolated in Panama for a prolonged period of time, followed by limited dispersal from this region to the rest of the Caribbean and Tropical Western Atlantic leading to its co-distribution. In this current study, we had to trade off increasing the number of samples sequenced per locality to increase the sampling distribution to include the entire range of the species. Increased sampling and sequencing for *B. annulata* from each locality throughout the region may be warranted and provide a better idea of the abundance of our Clade 1 lineage.

Many species with overlapping broad-scale biogeographic distributions can become specialized in different ecological niches and habitats, and thus, may become physically isolated from each other on a fine-scale (termed micro-allopatry; Bird et al., 2012; Getz & Kaitala, 1989; Tobler, Riesch, Tobler, Schulz-Mirbach, & Plath, 2009). For the *B. annulata* species complex, we found no obvious ecological or habitat difference that could explain a micro-allopatric divergence scenario. Samples from both putative lineages were collected from the same reef sites, at the same depths, and with the same crustacean symbionts. Our sample localities across the TWA span a continuum of reef environments from shallow, high-nutrient, nearshore patch reef sites in Bocas del Toro, Panama, to fore reefs along the Meso-American Barrier reef in Mexico, and Bahamian patch reefs surrounded by seagrass. Further, in many micro-allopatric divergence scenarios, habitat specialization is expected to lead to immediate isolation and a cessation of gene flow should follow (Bird et al., 2012; Tobler et al., 2009). Our model selection analyses prefer models with continuous unidirectional gene flow rather than models where a period of isolation followed divergence and that contemporary gene flow is the result of secondary contact. For these reasons, we conclude that the *B. annulata* species complex likely evolved in sympatry. Additional sampling with a greater focus on sampling across disparate habitats at each sample locality may provide greater clarity as to whether any micro-allopatric divergence scenarios are responsible for the divergence we have recovered.

In addition to overlapping biogeographic ranges, no obvious habitat partitioning, and a genomic signature of divergence with continuous gene flow, our genome scan analyses recover a number of loci that appear to be under selection (Table 6; Fig. S1). These include loci that are under putative divergent selection, and those that are more conserved relative to their neutrally evolving counterparts. The genomic underpinnings that may drive divergence and maintain reproductive isolation, even with continuous gene flow, are unknown in benthic anthozoans. In the stony coral genus *Acropora,* the *PaxC* gene (a nuclear intron) resolves a tree topology that clusters conspecifics that spawn in the same seasons, but no additional functional genes under putative natural selection have been identified (Rosser et al., 2017). A recent study also highlights the importance of variation in gene expression in maintaining ecological divergence (i.e. polygenic adaptation) across a sympatrically distributed coral species complex (Rose et al., 2018).

While our dataset is limited because only loci from *B. annulata* that map to the *E. pallida* genome are able to be identified, and we did not conduct a comparative transcriptomic investigation, our results may at least provide a basic starting point for future genomic investigations into tropical anthozoan speciation. All loci detected by BayeScan as being under divergent selection were mapped to the annotated *E. pallida* genome to identify loci that may be linked to functional genes. Five of the six loci were mapped to mRNA transcripts and CDS/exon regions (Table 6). Locus 1 mapped to a mRNA transcript linked to a predicted neuronal acetycholine receptor subunit alpha gene, a cell membrane receptor involved in muscle activation when acetylcholine is released by motor nerves in the central nervous system. Locus 97 mapped to a 3’ untranslated region (UTR) in an mRNA transcript linked to a predicted C-C chemokine receptor type 4. The C-C chemokine receptors are well characterized cell signaling pathways involved in immune responses (e.g. Murphy, 1994). The type 4 receptor transports leukocytes in vertebrates. Genes involved in vertebrate immune cell communication have been shown to evolve rapidly, thought to be a response to intense selective pressures placed on them by the molecular mimicry of microbes disrupting host immune responses (Bajoghil, 2013; Zlotnik, Yoshie, & Nomiyama, 2006). Other loci under selection that mapped to transcriptomic regions of the *E. pallida* genome include membrane proteins involved in lipid binding and recognition, vesicular trafficking, signal transduction, and phagocytosis (locus 197; den Hertog, 1999), a candidate toxin protein that is a well characterized component of snake venom (locus 1160; Doley & Kini, 2009), and proteins that bind to myosin and are involved in muscle contraction (Locus 1326). While it’s unclear whether, how, or why any of these identified functional genes maintain linkage disequilibrium, and thus putative species boundaries, ecological specialization could certainly drive adaptive divergence in immune response if different niches are exposed to different pathogens. It is also not hard to imagine that a shift in exposure to different prey types across disparate habitats could lead to a corresponding shift in venom composition or toxicity. The finding that 5/6 loci under divergent selection mapped to mRNA transcripts, and 2 of these loci mapped within CDS/exon regions is consistent with the idea that selection may play an important role in maintaining species boundaries between these cryptic taxa.

### 4.2. Tropical sea anemone diversity

*Bartholomea annulata* is the first tropical anemone species complex to be delimited using genomic data and a molecular systematic approach. With simple body plans, hydrostatic skeletons, no rigid structures, and few diagnostic characters (most of which are highly convergent; e.g. Rodriguez et al., 2014), morphological studies of sea anemones are challenging (Fautin, 1988), and much of the species-level diversity that exists could be cryptic. The lack of informative morphological characters is compounded by the history of molecular biology and the slow rate of mitochondrial genome evolution in anthozoans (Daly, Gusmão, Reft, & Rodríguez, 2010; Shearer, Van Oppen, Romano, & Wörheide, 2002). Short regions of mitochondrial DNA barcodes (mtDNA) became the molecular marker of choice for evolutionary biologists conducting population-level studies in most animal phyla (e.g. Avise, 2009). An unintended byproduct of these early studies was that biologists began uncovering highly divergent lineages in taxa that were nominally described as single morphological species. While anemone mtDNA is used in phylogenetics to resolve deeper taxonomic relatedness, it is incapable of picking up shallow divergence times, which are characteristic of many cryptic species complexes (Daly et al., 2010). Similarly, the universal molecular markers from the nuclear genome that are used for phylogenetic reconstruction are also too slowly evolving for population level studies (Daly et al., 2010). Finally, sea anemone diversity peaks in temperate regions where they are often the dominant benthic macrofauna, but lack species diversity in the tropics, which are dominated by scleractinian corals and octocorals (Fautin, Malarky & Soberon, 2002). This convergence of factors has led to a lack of evolutionary attention to tropical anemone species compared to temperate ones, and diversity in the tropics is likely under described as a result. Our findings demonstrate the utility of genomic data for discovering and delimiting cryptic anemone species that are likely to be missed using conventional markers. An important note, however, is that we used BFD* for our genomic species delimitation analyses. BFD* implements the multi-species coalescent model, which has been shown to be unable to distinguish between intraspecific population genetic structure and species boundaries under some speciation models (Sukumaran & Knowles, 2017). With this in mind, we suggest a targeted morphological study of both delimited lineages, and possibly an RNAseq approach to search for fixed SNPs and variation in expression levels in functional genes we have preliminarily identified here. Additional types of data could independent lines of evidence in support of our analyses here.

## Acknowledgements

We are grateful to Erich Bartels, Annelise del Rio, Jose Diaz, Dan Exton, Lisle Gibbs, Natalie Hamilton, Alex Hunter, Anna Klompen Jason Macrander, Spencer Palombit, Stephen Ratchford, Nuno Simoes, Jill Titus, Cory Walter, Eric Witt, Clay Vondriska, and the Operation Wallacea dive staff for assistance in the field and laboratory. We also thank Jordan Satler, Megan Smith, and Bryan Carstens for discussion and advice regarding *fastsicoal2* analyses and model selection. Bellairs Research Station, the Bermuda Institute of Ocean Science, Cape Eleuthera Institute, CARMABI, Coral View Dive Center, Gerace Research Centre, the Honduran Coral Reef Foundation, Mote Marine Laboratory, Smithsonian Tropical Research Institute, and the University of the Virgin Islands provided valuable logistical support in the field. Specimens were collected from throughout the Tropical Western Atlantic under permits: SE/A-88-15, PPF/DGOPA-127/14, CZ01/9/9, FKNMS-2012-155, SAL-12-1432A-SR, STT037-14, 140408, MAR/FIS/17, and 19985. This research was supported by funding from a National Science Foundation-Doctoral Dissertation Improvement Grant DEB-1601645 and Florida Fish and Wildlife Conservation Commission awards to B.M.T. & M.D. Operation Wallacea, American Philosophical Society, International Society for Reef Studies Graduate Fellowship, PADI Foundation Grant, and American Museum of Natural History Lerner Gray Funds funded field research for B.M.T. Additional funding was provided through the Trautman Fund of The OSU Museum of Biological Diversity, The Ohio State University, and National Science Foundation DEB-1257796 to MD.

## Data Accessibility Statement

Raw sequence data and all files for all analyses will be archived in Dryad upon final acceptance of this manuscript. Python scripts are also available on GitHub (github.com/pblischak/Bann_spdelim).

## Author Contributions

B.M.T. and M.D. conceived the study research; B.M.T. collected samples and conducted laboratory work. B.M.T., P.D.B. analyzed the data and conducted bioinformatics work. B.M.T., P.D.B., and M.D. wrote and edited the manuscript.

